# A systematic investigation of endothelial cell behavior under hydrostatic pressure

**DOI:** 10.1101/2025.05.27.656423

**Authors:** Giulia Venturini, Advik Sikligar, Giulia De Campo, David T. Eddington

## Abstract

Biomechanical stimuli are critical in regulating cell behavior and phenotype across various tissues and organs, particularly within the cardiovascular system. Endothelial cells, which line blood vessels, are continuously subjected to forces generated by the pulsatile nature of blood flow, including shear stress, strain, and hydrostatic pressure (HP). Among these stimuli, HP remains the least explored, primarily due to the technical challenges of incorporating it into conventional cell culture systems. However, HP significantly influences key biological processes, such as cell differentiation, migration, proliferation, and apoptosis. To facilitate the introduction of HP in vitro, we have previously developed an automated, high-throughput platform compatible with standard 96-well plates capable of delivering up to 12 independent pressure conditions. In this study, we applied this setup to investigate the effects of a wide range of static pressure conditions on the viability, morphology, and cytoskeleton adaptation of Human Umbilical Vein Endothelial Cells (HUVECs).

## 1. INTRODUCTION

The vascular endothelium lining the inner surface of blood vessels is constantly subjected to spatially and temporally changing biochemical and biomechanical forces. The pulsatile nature of blood flow in the elastic vasculature generates hemodynamic forces such as shear stress, hydrostatic pressure (HP), and cyclic strain, all of which influence endothelial cell (EC) behavior. Cardiovascular diseases such as hypertension and atherosclerosis are associated with disturbed hemodynamic patterns and complex multi-axial changes to the microenvironmental stress field, leading to altered EC function and morphology^1^. The imbalance of endothelial-derived factors resulting from endothelial dysfunction manifests as reduced vasodilation, increased inflammation and thrombosis, impaired angiogenesis, and heightened vascular permeability^2^. These phenotypic changes are driven by alterations in endothelial gene transcription mediated by mechanical forces^3^. For instance, genes involved in vascular tone regulation (NOS3, endothelin-1, and prostacyclin), thrombosis, and inflammation (platelet-derived growth factor, VCAM-1) contain shear stress response elements (SSREs) within their promoters regulating their transcription^4–6^. Additionally, the spatial and temporal dynamics of shear stress can trigger coordinated up- or down-regulation of clusters of genes relevant to vascular pathology^7^.

While the role of shear stress in endothelial dysfunction has been extensively studied, the impact of hydrostatic pressure (HP) on endothelial behavior remains comparatively underexplored. Previous studies suggest hydrostatic pressure is critical in vascular homeostasis and pathology^8^. Chronic exposure to HP (2.9–150 mmHg) has been shown to enhance proliferation and promote multilayering in aortic ECs^9^. When exposed to aortic pressure (50–150 mmHg), these cells exhibit three-dimensional cytoskeletal rearrangements and time-dependent morphological changes^10,11^. In contrast, HUVECs do not appear to undergo morphological adaptation in response to aortic HP but instead display increased contractility mediated by VE-cadherin reorganization and actin stress fiber formation^12–14^. Notably, research on HUVECs mostly focuses on supraphysiological pressure levels (50, 100 mmHg) while largely overlooking physiologically relevant ranges (10–30 mmHg). Few studies have examined the effects of venous pressure, reporting cell elongation under 25 mmHg^11^ and increased proliferation under cyclic pressure (60/20 mmHg)^15^. Recently, capillary HP (20 mmHg), but not arterial pressure (55 mmHg), has been shown to promote sprouting angiogenesis in HUVECs via YAP (yes-associated protein I) signaling^16^, further emphasizing the need to investigate endothelial behavior under pressure conditions that better reflect *in vivo* physiology.

The lack of research on HP-mediated mechanotransduction is primarily due to technical challenges in incorporating hydrostatic pressure in standard *in vitro* systems and isolating its effects from shear stress. Current approaches, such as syringe pumps^12,14,17^, media column height^16,18,19^, or gas pressurization^13,20^, have been used to apply HP *in vitro*, but they suffer from key limitations: (1) they support only one pressure condition at a time; (2) they are incompatible with high-throughput analysis tools like plate readers; and (3) when dynamic pressure waveforms are implemented, they are typically limited to simplified profiles such as sinusoidal, triangular, or square waves. To address these constraints, we have previously developed a high-throughput, automated device compatible with standard cell culture platforms, such as the 96-well plate^21^. Our system enables the application of up to 12 independent pressure waveforms with fully customizable magnitude, frequency, and duration. However, in the current study, we are limiting to static pressure magnitudes; dynamic pressure waveforms will be explored in future work. Here, we leverage the high-throughput capabilities of our platform to systematically investigate HUVECs behavior across a wide range of physiologically and pathologically relevant hydrostatic pressures (10, 20, 50, 75, 100, and 150 mmHg). We characterize pressure-dependent changes in morphology and cytoskeletal organization and examine the impact of HP on cell viability, redox activity, and monolayer integrity.

## 2. EXPERIMENTAL SETUP AND METHODS

### 2.1. Pressure setup construction

The setup used in this study was previously described^21^. Briefly, the system can deliver up to 12 different pressure conditions to cells cultured in a 96-well plate. The pressure in each line is controlled by a proportional valve (Parker VSO LowPro) and measured by a pressure sensor (Honeywell 40PC series), both interfacing with an Arduino Nano microcontroller. The Arduino monitors the pressure in real-time and adjusts the valve aperture via Pulse Width Modulation (PWM) through a PID control loop to maintain the desired setpoint. Pressurized gas (5% CO_2_, 21% O_2_, balanced N_2_), supplied by a gas cylinder, is distributed to the headspace of each well of a plate’s column through a millifluidic 3D-printed insert (Fig. 1a,1b). Tight sealing of the plate is ensured by a set of three O-rings (one Buna-N O-ring and two PFTE backup rings) mounted on each column of the device (Fig. 1c,1d).

**Figure 1:**
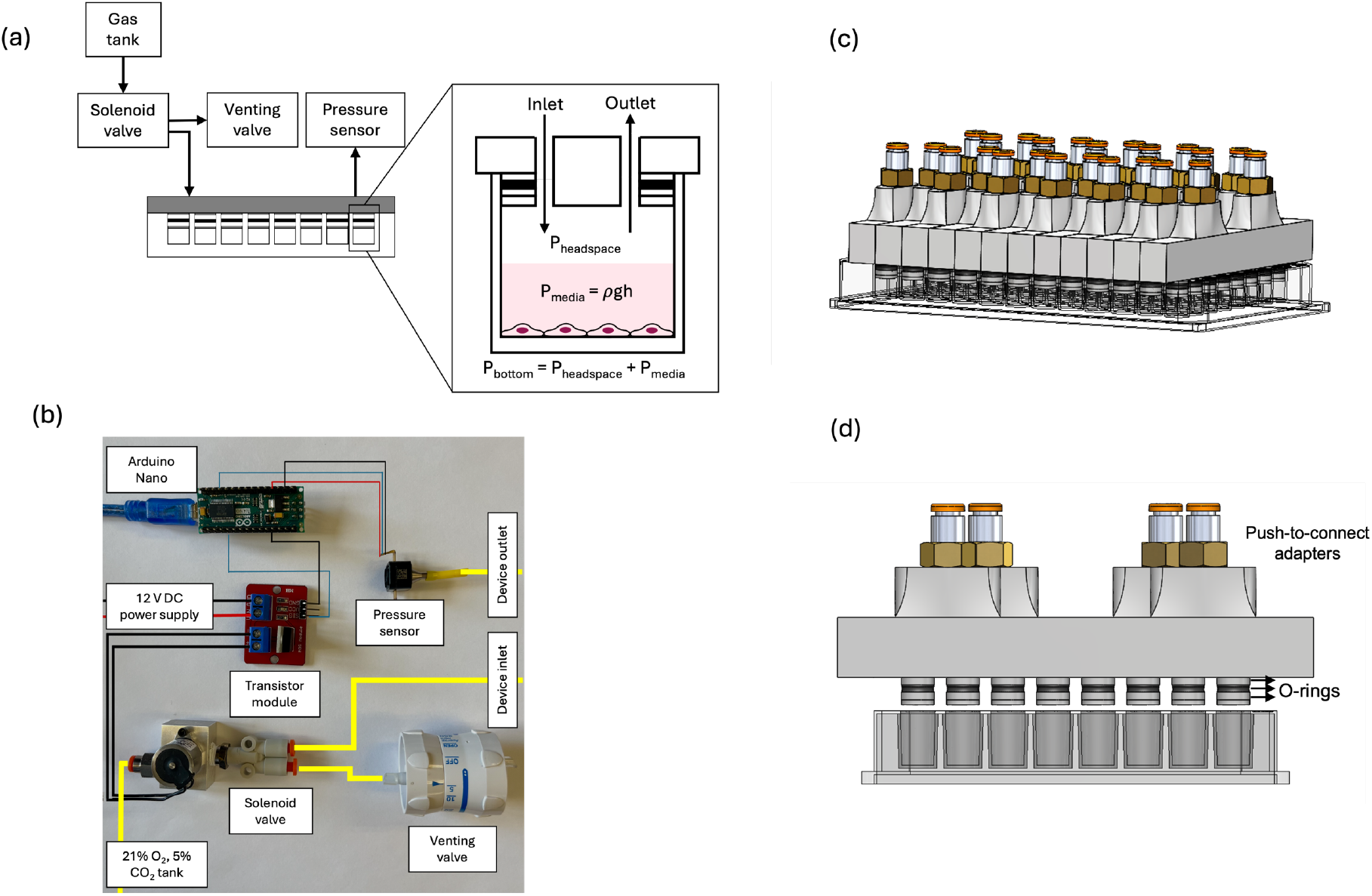
Pressure delivery system. a) Schematic showing pressure control in one column of the plate and in a single well. b) Components needed for pressure control in one column of the plate. c) 3D CAD of the device inserted in the plate. d) 3D CAD of the front-view of the device showing O-rings and connectors.

### 2.2. Cells

Human umbilical vein endothelial cells (Lonza) were grown in EBM-2 medium supplemented with EGM-2 bullet kit (Lonza) and 1% penicillin streptomycin (Gibco BRL) at 37°C and 21% O_2_ and 5% CO_2_. Cells were split at 80% confluence, and passages 2-8 were used for the experiments.

### 2.3. Pressure treatment

Before each experiment, the wells of a 96-well plate were coated with a 2% gelatin solution (Sigma Aldrich). Cells were then seeded at a density of 15,625 cells/cm^2^ and allowed to grow for 24 hours. Subconfluent monolayers were exposed to various static pressure conditions (10, 20, 50, 75, 100, 150 mmHg above atmospheric pressure) for 36 hours. Control cells were grown at ambient pressure. To isolate the contribution of hydrostatic pressure from other mechanical forces in the hemodynamic milieu (shear stress and stretch), cells were cultured in no-flow conditions on a rigid substrate, and the gas phase above the supernatant was pressurized, imposing a normal force equal to the applied pressure.

### 2.4. Cell viability and redox assay

Viability was measured using live/dead staining. Briefly, cells were stained with 1 μg/ml of propidium iodide (PI) and 5 μg/ml of Hoechst 33342. Cells were incubated in the dark for 15 minutes at room temperature and imaged to obtain the total number (Hoechst) and the number of dead cells (PI). Cell redox activity was assessed using PrestoBlue Reagent (ThermoFisher), according to manufacturer instructions. The amount of fluorescent resorufin resulting from resazurin reduction was read with a plate reader (Varioskan, Thermo Scientific) at 560/590 nm excitation/emissions (bottom read) and normalized by the total number of cells in each well.

### 2.5. Staining

After pressure exposure, cells were fixed and stained for morphology assessment. First, cells were incubated for 10 minutes at room temperature with a 4% paraformaldehyde in PBS (phosphate-buffered saline) solution. Cells were then permeabilized with a solution of 0.1% Triton-X in PBS for 10 minutes at room temperature and incubated overnight at 4°C with CD144 (VE-cadherin) monoclonal primary antibody (Invitrogen), diluted in a ratio of 1:200 in PBS, 1% BSA and 0.1% Tween-20. Cells were incubated for one hour at room temperature with Donkey anti-mouse IgG secondary antibody AlexaFluor 488 (Invitrogen) and AlexaFluor 647 phalloidin (Invitrogen, F-actin stain) at a ratio of 1:500 and 1:100, respectively, in PBS, 1% BSA and 0.1% Tween-20. In the last 15 minutes of incubation, Hoechst 33342 was added to each well at a 5 μg/ml concentration to stain the cell nuclei.

### 2.6. Cell density, morphology, cytoskeleton remodeling, and cell junction analysis

After each experiment, fluorescence images were acquired for each condition at arbitrary locations in four wells at 40X and 10X magnification with an Olympus IX71 microscope. The pictures were post-processed with Fiji (ImageJ), and a total of 300 cells were analyzed for each condition. Cell density in 1 mm^2^ was estimated by automatically counting cell nuclei in 10X binary pictures using Fiji. Cell contours in 40X pictures were manually delineated for morphological assessment following the VE-cadherin signal. Cell area, circularity, and orientation angle were evaluated through Fiji. Cell tortuosity was instead calculated as the ratio between the cell perimeter and the perimeter of the equivalent ellipse. Nuclei area and aspect ratio were obtained through Fiji: binary images of the nuclei were generated, adjacent nuclei were separated using the “watershed” command, and morphological parameters were automatically derived. For cytoskeleton remodeling analysis, pictures were acquired at 20X magnification with a confocal microscope (Olympus FV3000). The cytoskeleton order parameter, S = <cos2ϑ>, was calculated for 30 cells in each condition to determine the degree of F-actin filament alignment to the major axis of the cell. For cell junction analysis, 30 cells for each condition were analyzed in 20X fluorescent pictures using Fiji to obtain VE-cadherin thickness and intensity profiles.

### 2.7. Statistics and reproducibility

Statistical analysis was performed using GraphPad Prism 10.4.2 software. ANOVA analysis with post-hoc Tukey’s test was conducted to compare groups that were normally distributed. Unless stated elsewhere, all experiments were performed with at least three biological independent replicates. Statistical significance was set as follows: *p < 0.05, **p < 0.01, ***p < 0.001****p < 0.0001. Statistical tests and relative p values are indicated in each figure legend.

## 3. RESULTS

### 3.1. Cell viability, density, and mitochondrial activity

HUVECs were cultured for 36 hours under static hydrostatic pressure ranging from 10 to 150 mmHg. Cells in all conditions showed similar viability to the control (Fig. 1a). Cells cultured under HP displayed a lower density than the control, correlated to the magnitude of applied pressure (Fig. 2b). The redox assay (Fig. 2c) showed overall higher reducing function in cells treated under pressure.

**Figure 2:**
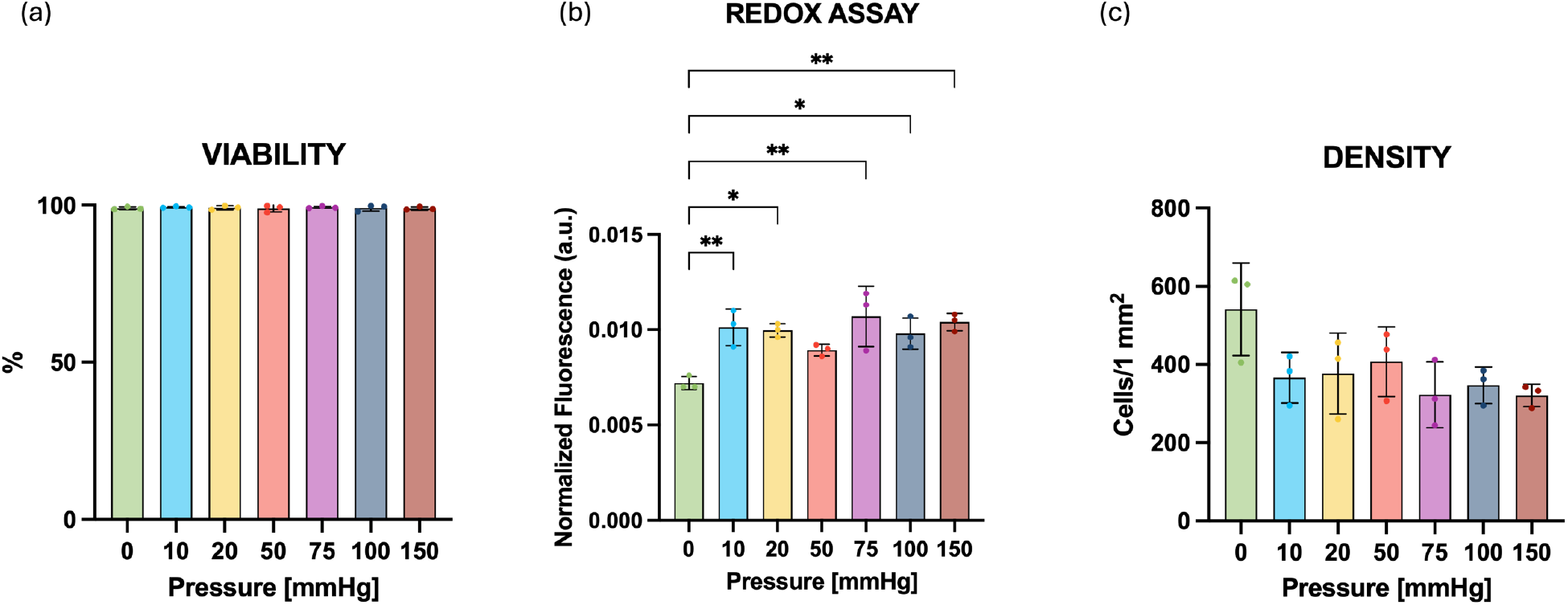
Cell viability, density, and redox activity under HP. a) Quantification of cell viability after 36h under pressure. The graph shows mean + s.d. N=3 independent experiments. b) Quantification of cell metabolic function after 36h under pressure. The graph shows mean + s.d. N=3 independent experiments. One-way ANOVA with Tukey’s post hoc test. *p<0.05, **p<0.01. c) Quantification of cell density after 36h under pressure. The graph shows mean + s.d. N=3 independent experiments.

### 3.2. Cell morphology

After pressure conditioning, cells were stained and imaged for morphological assessment. Hydrostatic pressure induced significant elongation (Fig. 3a), with cells displaying lower circularity than those cultured at ambient pressure. The elongation was most pronounced under pressures resembling venous levels but diminished at higher magnitudes. Additionally, cell area increased with pressure (Fig. 3b), suggesting enhanced spreading onto the substrate. Similar trends were observed in cell nuclei, which exhibited slight increases in both area and aspect ratio, consistent with the elongation and spreading of the cells (Fig. 4a-c). Across all conditions, cells appeared randomly oriented (Fig. 3c) with comparable tortuosity (Fig. 3d).

**Figure 3:**
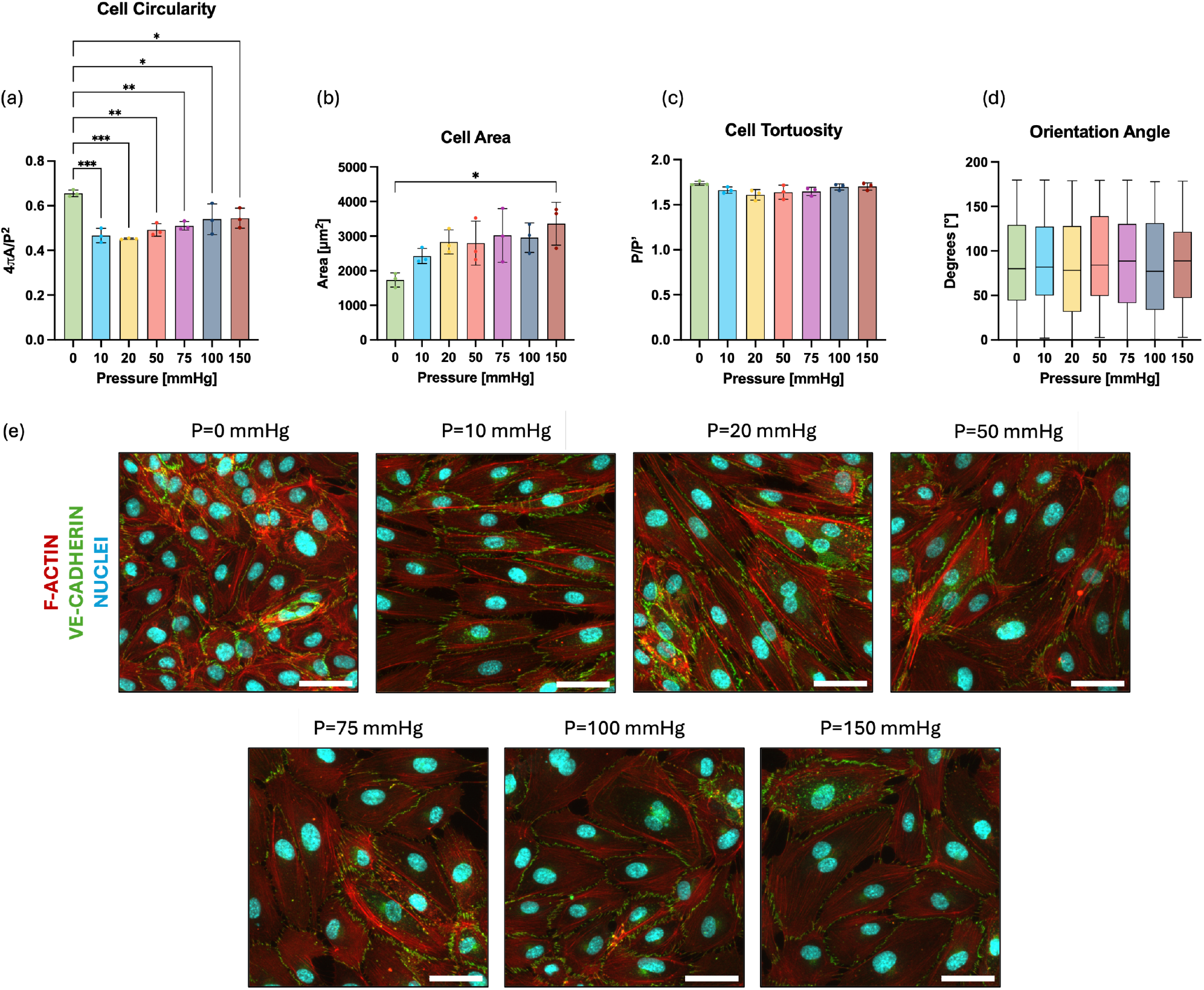
Cell morphological adaptations to HP. a) Quantification of cell circularity after 36h under pressure. The graph shows mean + s.d. N=3 independent experiments, n≥ 250 cells analyzed per group. One-way ANOVA with Tukey’s post hoc test. *p<0.05, **p<0.01, ***p<0.001. b) Quantification of cell area after 36h under pressure. The graph shows mean + s.d. N=3 independent experiments, n≥ 250 cells analyzed per group. One-way ANOVA with Tukey’s post hoc test. *p<0.05. c) Quantification of cell tortuosity after 36h under pressure. The graph shows mean + s.d. N=3 independent experiments, n≥ 250 cells analyzed per group. d) Cell orientation angle distribution after 36h under pressure. The graph shows the median and 25th to 75th percentile; whiskers indicate min and max values. N=3 independent experiments, n≥ 250 cells analyzed per group. e) Representative confocal z-projection images of HUVECs showing VE-cadherin (green), F-actin (red), and nuclei (cyan) after 36h pressure exposure. Scale bar: 50 μm.

**Figure 4:**
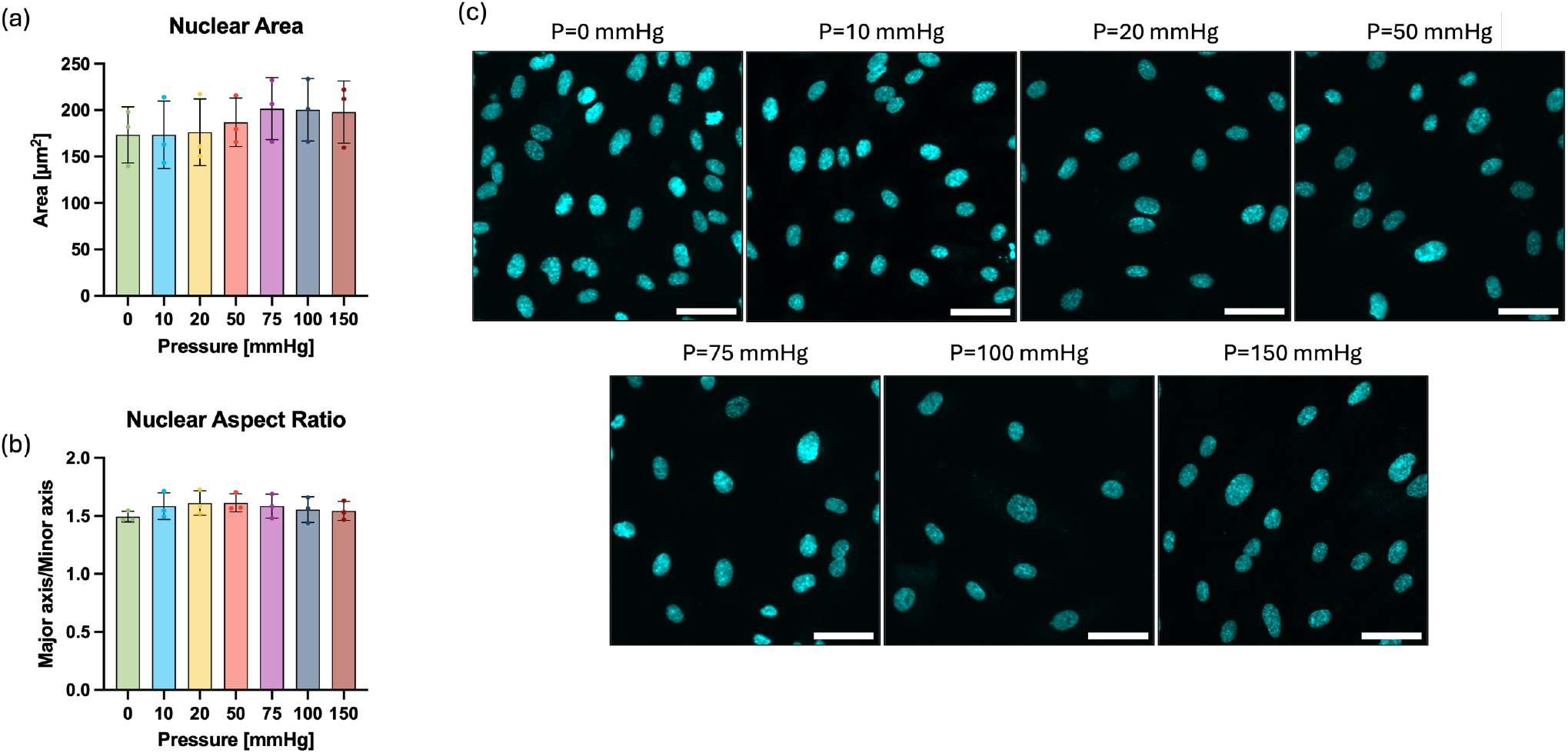
Nuclear morphological changes under HP. a) Measurement of nuclei area after 36 hours of pressure treatment. The graph shows mean + s.d. N=3 independent experiments, n≥ 200 cells analyzed per group. b) Quantification of nuclei aspect ratio after 36 hours under pressure. The graph shows mean + s.d. N=3 independent experiments, n≥ 200 cells analyzed per group. c) Representative confocal z-projection images of nuclei after 36h pressure exposure. Scale bar: 50 μm.

### 3.3. F-actin and cytoskeleton remodeling

F-actin staining (Fig. 5b) revealed marked cytoskeletal reorganization in response to hydrostatic pressure. Cells treated under venous pressure (10, 20 mmHg) displayed parallel stress fibers aligned with the major axis and distributed throughout the cytoplasm. Fiber alignment progressively decreased at increasing pressure magnitudes (50-150 mmHg). In contrast, cells cultured at ambient pressure exhibited randomly oriented F-actin fibers localized predominantly at the periphery. Quantitative analysis of F-actin alignment (Fig. 5a-c) confirmed these observations, with cytoskeletal parameter values ranging from 0.68 (150 mmHg) to 0.95 (20 mmHg) under pressure, indicating a higher degree of alignment, compared to 0.45 in the control condition.

**Figure 5:**
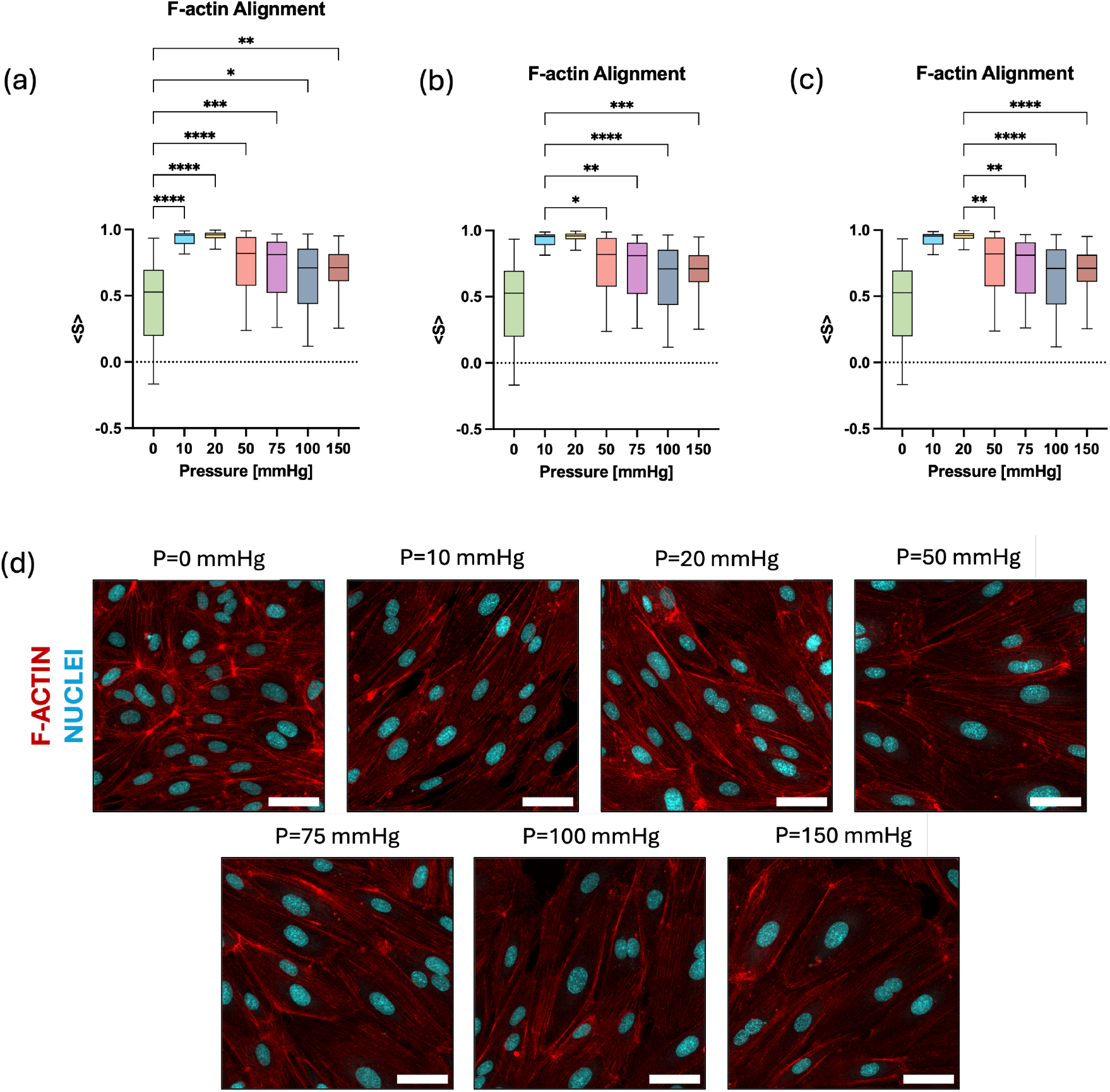
Cytoskeleton remodeling under HP. a) Quantification of cell cytoskeleton order parameter, <S>, after 36h under pressure. The graph shows the median and 25th to 75th percentile, whiskers indicate min and max values; N=3 independent experiments, n≥ 30 cells analyzed per group. One-way ANOVA with Tukey’s post hoc test, comparison between control and pressure conditions. *p>0.05, **p>0.01, ***p>0.001, ****p>0.0001. b) Quantification of cell cytoskeleton order parameter, <S>, after 36h under pressure. One-way ANOVA with Tukey’s post hoc test, comparison between 10 mmHg and the other pressure conditions. *p>0.05, **p>0.01, ***p>0.001, ****p>0.0001. c) Quantification of cell cytoskeleton order parameter, <S>, after 36h under pressure. One-way ANOVA with Tukey’s post hoc test, comparison between 20 mmHg and the other pressure conditions. *p>0.05, **p>0.01, ***p>0.001, ****p>0.0001. d) Representative confocal z-projection images of F-actin fibers and nuclei in HUVECs after 36h pressure exposure. Scale bar: 30 μm.

### 3.4. VE-cadherin and monolayer integrity

VE-cadherin staining (Fig. 6a) in cells cultured at ambient pressures showed continuous, mature cell-cell junctions (orange arrows) characterized by thick actin bundles parallelly aligned and not overlapping with VE-cadherin at the junction. Venous pressure conditions (10, 20 mmHg) showed thicker, continuous junctions along the cell side and VE-cadherin/actin protrusions (white arrows) at the apical and basal sides. Cells cultured at 50 and 75 mmHg pressures showed continuous, although thinner, junctions and reduced VE-cadherin/actin protrusions. At supraphysiological levels (100, 150 mmHg), junctions became more fragmented and discontinuous (blue arrows). We extracted normalized intensity profiles across cell-cell contacts to assess VE-cadherin expression and width of endothelial junctions. As shown in Fig. 6b, when exposed to venous pressure (10, 20 mmHg), cells exhibited higher fluorescent signal and wider intercellular junctions. The expression of VE-cadherin and width of cell junctions progressively decreased at higher pressure magnitudes, going below control levels at 100 and 150 mmHg.

**Figure 6:**
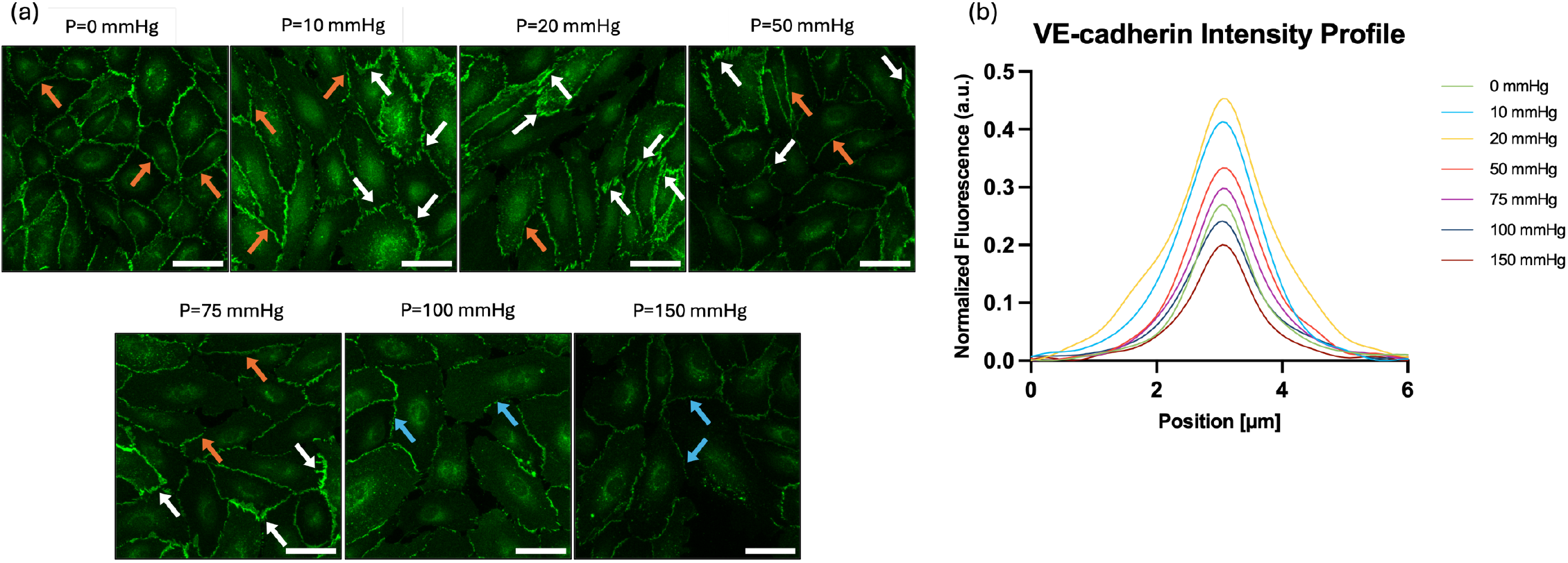
HP effect on monolayer integrity. a) Representative confocal z-projection images of VE-cadherin in HUVECs after 36h pressure exposure. Scale bar: 50 μm. Orange arrows indicate continuous cell-cell junctions, white arrows point to VE-cadherin/actin protrusions, and blue arrows indicate fragmented junctions. b) Normalized mean intensity profiles of VE-cadherin junctions after 36 hours of exposure. N=3 independent experiments, n≥ 30 cells analyzed per group.

## 4. DISCUSSION

Research on hydrostatic pressure’s effect on endothelial cell behavior is limited, yet previous studies have shown its crucial role in cell proliferation^9,22,23^, apoptosis^15,24^, vascular angiogenesis^16,25,26^, and monolayer integrity^19,27^. A major challenge in studying HP-mediated mechanotransduction is implementing hydrostatic pressure *in vitro* while isolating its effects from shear stress. Current methods introducing HP in cell cultures are limited to a single pressure condition at a time, significantly reducing the experimental throughput. Studies on venous endothelial cells have predominantly focused on supraphysiological pressures (>50 mmHg), with relatively few investigating physiologically relevant venous ranges. Notably, HUVECs behavior appears to be pressure-dependent, with phenotypic changes observed primarily under venous pressures^11,15,16^. However, the substantial variability in experimental setups and parameters complicates direct comparisons, with contrasting evidence on cell response based on pressure magnitude and frequency^11,12,14,15^. To address these discrepancies, we systematically investigated HUVECs response across various physiologically relevant pressure conditions (10, 20, 50, 75, 100, 150 mmHg). Our high-throughput system enables the simultaneous application of up to 12 independent, fully customizable pressure waveforms in standard 96-well plates, significantly reducing experimental time and facilitating downstream analyses, such as plate reader-based assays. However, in this study, we limited to six static pressure conditions. Leveraging the system’s capabilities, we examined the impact of HP on cell viability, redox activity, morphological and cytoskeletal adaptations, and junctional remodeling.

Consistently with our previous findings^21^, cell viability under pressure remained comparable to atmospheric pressure conditions, even at supraphysiological levels, as shown in Fig. 2a. Although cells proliferated under pressure, the final density after 36 hours was lower at higher pressures (Fig. 2c). At the same time, cells under increasing pressure exhibited a larger surface area (Fig. 3b). This apparent decrease in density is unlikely to result from pressure-induced cell death or detachment, as neither was observed during the experiments. Instead, larger individual cell areas may reduce the available space for proliferation, leading to contact inhibition and a lower overall cell number.

Cells cultured under HP showed substantial morphological changes. In agreement with previous findings^11^, we observed random cell elongation under venous pressure, which reduced at supraphysiological pressure levels, as shown in Fig. 3a. Structural changes at the cellular level were reflected in the cell nuclei, even if to a much smaller degree (Fig. 4). The observed morphological adaptation results from mechanotransduction processes mediated by the actin network, enabling cells to optimize their shape and enhance structural integrity in response to mechanical stress. Indeed, a recent study showed that early cell response to pressure is primarily driven by the cytoskeleton, with mechanical loading triggering an actomyosin-mediated cell contraction already after 5 minutes of exposure^13,14^. Similarly to previous studies^10,28^, our results (Fig. 5) show significant actin reorganization under pressure, particularly under venous pressure magnitudes, with parallel stress fibers evenly distributed throughout the cytoplasm and aligned along the major cell axis. Cells cultured at ambient pressure instead exhibited randomly oriented F-actin filaments localized predominantly at the periphery. Alongside cell elongation, actin cytoskeletal reorganization resulted in cells adopting larger surface areas at higher pressure magnitudes (Fig 3b), which has previously been shown to increase cell adhesion to the underlying substrate and provide mechanical stability^29^. Although not explored in this work, such spreading is likely driven by cell focal adhesion remodeling prompted by actomyosin cytoskeleton contraction and mediated by integrins^9,30,31^.

Cells under pressure exhibited increased redox activity, as shown in Fig. 2b. The primary sources of reducing power are mitochondria and NADPH oxidases. Mitochondrial activity is closely associated with intracellular Ca^2+^ levels^33^, which rapidly increase upon pressure exposure as part of the mechanotransduction process mediated by mechanosensitive cation channels^13^. This Ca^2+^ influx initially triggers actomyosin contraction through myosin light chain phosphorylation^30^. While the long-term dynamics of Ca^2+^ signaling under chronic pressure exposure remain unclear, mitochondria could participate in buffering intracellular Ca^2+^. HUVECs show a shear stress-induced Ca^2+^ response caused by the activation of the PLC/IP_3_/IP_3_R pathway, with mitochondria exchanging Ca^2+^ with the endoplasmic reticulum to coordinate intracellular Ca^2+^ transients and oscillations^34^. A similar IP_3_-evoked Ca^2+^ release has been observed in pressurized arteries^35^, suggesting that mitochondria could also participate in Ca^2+^ signaling during HP response. Mitochondrial control of Ca^2+^ signaling has been shown to be mediated by ATP production^36^ and to involve the generation of reactive oxygen species (ROS) as a protective mechanism against Ca^2+^ overload during IP_3_-mediated release^37^. This buffering mechanism could contribute to the increased redox activity observed under pressure. ROS generation under HP may also modulate cytoskeletal remodeling. For instance, hypertension is known to contribute to NADPH oxidases-mediated vascular oxidative stress and endothelial dysfunction through the activation of the ILK-1/βPIX/Rac-1 signaling pathway^38^, suggesting an interplay between integrin-linked cytoskeletal signaling and cellular redox regulation. Moderate amounts of cytosolic and mitochondrial ROS have been shown to regulate actin cytoskeleton organization during cell migration^39^, microvascular remodeling^40^, wound healing^41^, and post-hypoxic recovery^42^, and might similarly contribute to cellular adaptation under hydrostatic pressure, potentially explaining the elevated redox activity observed in our experiments. Nonetheless, further investigation is needed to fully elucidate the redox mechanisms triggered by pressure and the cross-talk between ROS and cytoskeletal dynamics.

Exposure of endothelial monolayers to hydrostatic pressure caused a marked redistribution of VE-cadherin at cell-cell junctions, as shown in Fig. 6. At venous pressure levels (10, 20 mmHg), cells displayed broader junctions and VE-cadherin/actin-rich protrusions, indicative of active cytoskeletal remodeling. Junctional strengthening under these conditions is likely supported by the formation of actin-driven membrane structures, such as junction-associated lamellipodia, which have been shown to dynamically reinforce adherens junctions by promoting cadherin clustering^43^. The observed protrusions resemble actin-related protein 2/3 complex (ARP2/3)-dependent extensions like cadherin fingers, previously described as elements orchestrating structural organization in migrating cells^44^. In our context, these structures may play a similar role by coordinating morphological adaptations between neighboring cells. As pressure increased to intermediate levels (50, 75 mmHg), VE-cadherin staining remained continuous but appeared thinner, with less actin-rich protrusions. At supraphysiological pressures (100, 150 mmHg), VE-cadherin localization became fragmented and discontinuous, indicating junction destabilization. These observations align with previous results suggesting that junctional stability depends on a three-way feedback between active force generation, cadherin bond turnover, and actin polymerization^45^. In this framework, mechanical cues such as hydrostatic pressure activate Src kinase, which initiates downstream signaling by activating Rho GTPases, specifically RhoA and Rac-1. Changes in the mechanical microenvironment can affect the interplay between these two factors, either strengthening or disrupting cell junction integrity. RhoA activation enhances actomyosin contractility, generating junctional tension that supports cadherin clustering, while Rac-1 promotes cell-cell contact through membrane protrusions by activating the WAVE complex regulator responsible for ARP2/3-mediated actin polymerization. Excessive RhoA activation can increase actomyosin tension, leading to bond disassociation, while low Rac1 activity impairs actin protrusion and intercellular gap closure, resulting in compromised monolayer integrity. Our results reflect this mechanistic balance: physiological pressure promotes adaptive remodeling and adhesion reinforcement, whereas higher pressures disrupt signaling homeostasis, driving junction disassembly.

In summary, we showed that HUVECs respond to hydrostatic pressure in a magnitude-dependent manner. Specifically, physiological venous HP (10, 20 mmHg) induced marked morphological and cytoskeletal adaptations and supported monolayer integrity by strengthening cell-cell junctions. Although higher pressure values also triggered structural rearrangements, their effects were less pronounced, and chronic exposure to supraphysiological HP significantly disrupted VE-cadherin expression and junctional stability. HP significantly affected cell metabolic activity, suggesting an interplay between cytoskeletal adaptations and redox regulation. Altogether, our findings highlight the importance of introducing HP as an experimental variable to develop more physiologically relevant *in vitro* models of the endothelium. Future studies will combine live-cell imaging of intracellular Ca^2+^ levels and quantification of ROS production to dissect the spatiotemporal interplay between Ca^2+^ signaling, redox activity, and cytoskeletal remodeling under dynamic, physiologically relevant hydrostatic pressure waveforms.

## 5. CONCLUSION

In this work, we sought to systematically investigate HUVECs behavior under a wide range of hydrostatic pressure values by leveraging the high-throughput nature of our pressure-control system. Our findings demonstrate that HUVECs undergo pressure-dependent morphological and metabolic changes, primarily linked to actin cytoskeleton remodeling. We also observed distinct changes in VE-cadherin junctions, with enhanced barrier integrity at physiological venous pressures and progressively thinner, fragmented cell-cell connections at higher pressures. Overall, our systematic approach might enable a deeper understanding of the phenotypic adaptations of HUVECs to both physiological and supraphysiological pressures, gaining valuable insights into the role played by HP in endothelial dysfunction. The high-throughput capabilities of our system and its compatibility with conventional culture platforms could pave the way to more efficient and standardized studies on endothelial behavior under pressure.

## 8. ACKNOWLEDGMENTS

The authors acknowledge Dr. Salman Khetani for generously granting access to the confocal microscope in his laboratory.

## 9. AUTHORS CONTRIBUTION

D.T.E. conceptualized the research, directed and supervised the study. G.V. designed and built the experimental setup, collected data, performed results analysis, and wrote the paper. A.S. and G.D. contributed to the development of the pressure system used in the study.

## 11. DATA AVAILABILITY

The data that support the findings of this study are available from the corresponding author upon reasonable request.

